# Differential regulation of myeloid Scavenger Receptor A (CD204) expression in humans versus mice in response to sepsis

**DOI:** 10.64898/2026.11.10.765736

**Authors:** Tammy R. Ozment, Melanie D. McGlothlin, James Adkins, Nikolas Hopkins, John B. Schweitzer, David L. Williams

## Abstract

Sepsis is the number one cause of mortality in the critically ill patient. The underlying pathophysiology of sepsis is not fully understood. Scavenger Receptor A (SR-A) is a promiscuous receptor of myeloid cells that binds multiple pathogen associated molecular patterns (PAMPs). We have demonstrated that SR-A promotes mortality in murine sepsis. The present study sought to examine the effect of sepsis on myeloid SR-A expression in septic mice and compare it to sepsis patients. Leukocytes were isolated from mice after cecal ligation and puncture. Blood leukocytes were isolated from sepsis patients. Myeloid cells were isolated from healthy mice or human subjects and were treated with PAMPs. Leukocyte phenotype and SR-A expression were determined by flow cytometry. CLP increased SR-A expression when compared to sham operated animals on both neutrophils and monocytes/macrophages. In contrast, SR-A expression was not changed on either neutrophils or monocytes from sepsis patients when compared to healthy control subjects. Treatment of murine neutrophils and macrophages with Pam3CSK4 and treatment of macrophages with LPS increased SR-A expression. Human neutrophil SR-A expression was not altered by any PAMPs, and only treatment with *C. albicans* glucan increased SR-A expression on human monocytes. Myeloid SR-A expression is increased in septic mice, and this increase plays a detrimental role in the response to sepsis, while SR-A expression is not changed in leukocytes from septic patients and does not correlate with SOFA II scores. Thus, myeloid SR-A expression does not appear to play significant a role in the pathophysiology of human sepsis.

**Summary Sentence:** SR-A (CD204) is differentially regulated in mouse versus human sepsis.

## Introduction

Sepsis is a life-threatening organ dysfunction in response to a dysregulated immune response to infection (1). It is estimated that there are ∼48.9 million cases of sepsis globally with 11 million sepsis related deaths (2). The mortality rate of sepsis patients correlates with disease severity and ranges from 25% for severe sepsis to 50% for septic shock (3, 4). Survival rates for sepsis have been improving due to improvements in diagnosis and supportive care, but attempts at developing specific therapies for sepsis have proven to be exceedingly difficult. This is due to our limited understanding of the factors that create the dysfunctional immune response to infection that leads to sepsis and sepsis syndrome. A better understanding of the molecular mechanisms that mediate immune dysfunction and septic sequelae may lead to advances that will improve outcomes.

Excessive inflammation with concurrent immunosuppression is a hallmark of sepsis, and this is mediated in part by dysfunctional immune cells (1). Septic neutrophils exhibit prolonged lifespan, defective microbial uptake and killing, and loss of targeted migration into sites of infection (5–7). This leads to release of neutrophil granular contents and neutrophil extracellular traps (NETs) in the microvasculature resulting in inflammation and organ damage (5–8). In sepsis, monocytes express lower levels of HLA-DR and higher levels of co-inhibitory molecules (9,10,12). These monocytes are unable to stimulate T cells and instead induce lymphocyte anergy or apoptosis, and ultimately, immunosuppression (9). Whether sepsis mortality is mediated by excessive inflammation or immunosuppression is controversial, but a better understanding of what drives the leukocyte dysfunction in sepsis could lead to new insights and the identification of novel therapeutic targets.

Scavenger Receptor A (SR-A, aka CD204) is a multi-functional receptor expressed by macrophages as well as neutrophils and dendritic cells (13–19). SR-A binds endogenous ligands including modified LDL and apoptotic cells (13,16,20,21) and pathogen associated molecular patterns (PAMPs) including endotoxin, lipoteichoic acid, and fungal glucans (16,20–24). SR-A’s role in atherosclerosis has been intensively studied (13,14), but more recently SR-A has been identified as a pattern recognition receptor in the innate immune system (15,16, 20, 25–29). The role of SR-A in response to infection is complex and appears to be greatly dependent upon the infectious agent. Though SR-A has been reported to be protective in several single organism infections (14,27,30), we (31) and others (32) have reported that SR-A deficient mice show improved survival, decreased inflammation, and decreased bacterial burdens in polymicrobial sepsis when compared to wild type mice (31). This suggests that SR-A might be a potential therapeutic target in sepsis.

The goal of the present study was to compare and contrast SR-A expression in septic mice and sepsis patients; and elucidate the effect of pathogen associated molecular patterns (PAMPs), often found in sepsis, on myeloid cell SR-A expression *in vitro*. Our goal was to determine whether SR-A may be a potential therapeutic target and/or biomarker for human sepsis.

## Materials and Methods

### Mice

ICR/HSD male mice were purchased from Envigo (Indianapolis, IN). The animals were maintained on standard laboratory chow and water *ad libitum* with a 12-hour light/dark cycle. Serologic testing confirmed that the mice were virus free. All animal procedures were reviewed and approved by the Institutional Animal Care and Use Committee at the James H. Quillen College of Medicine, East Tennessee State University.

### Patient samples

Healthy control blood (181 samples) was purchased from a licensed vendor (BioIVT, Johnson City, TN). After informed consent, whole blood was drawn from sepsis patients (105 samples) identified by clinicians on the Infectious Disease service who calculated Sequential Organ Failure Assessment (SOFA) II scores. All procedures were reviewed and approved by the East Tennessee State University Institutional Review Board.

### Polymicrobial Sepsis

Age and weight matched adult (3-6 months old) male mice underwent cecal ligation and puncture (CLP) as previously described to induce polymicrobial sepsis (34). Briefly, the cecum was exteriorized, ligated just distal to the ileocecal junction, and double punctured with a 20-gauge needle. Sham surgery (laparotomy alone) mice were used as a control for surgery and anesthesia, and animals that underwent no surgery or anesthesia were employed as negative controls. One ml of warmed lactated ringers was given IP for post-operative recovery. This model has an approximately 60% mortality rate at 7d (data not shown). Mice were sacrificed at 3, 16, or 24 h or 2 or 7d post-operatively.

### Murine Leukocyte Isolation and Cell Culture

ICR/HSD male mice were injected with 2 ml 4% thioglycollate IP. Mice were euthanized by CO_2_ asphyxiation 72 h later, and peritoneal cells were harvested by lavage. Alternatively, control animals were euthanized by CO_2_ asphyxiation and both femurs, tibias, and humeri were harvested. The marrow cavities were flushed with ice cold PBS to collect bone marrow cells and RBC were lysed hypotonically. Cells were cultured into 6 well plates in high glucose Iscove’s Modified Dulbecco’s Medium (IMDM, ATCC) supplemented with 10% heat inactivated horse serum, GM-CSF and antibiotics (bone marrow) or DMEM supplemented with 10% fetal bovine serum and antibiotics (peritoneal macrophages) and treated with Pam3CSK4, (10 μg/ml, InVivogen), ultrapure LPS (100 ng/ml, InVivogen), a combination of Pam3CSK4 and LPS, particulate glucan (10 μg/ml) or media alone in triplicate for 24 h.

### Human Leukocyte Isolation and Cell Culture

Neutrophils and Monocytes were isolated by negative magnetic separation using the EasySep Direct Human Neutrophil Isolation kit or EasySep Direct Human Monocyte Isolation kit, respectively (StemCell Technologies, Cambridge, MA) from 9 healthy subjects (3 males, 6 females; age range 27-48 years). Cells were counted using the Countess Cell Counter (ThermoFisher).

Trypan Blue staining indicated live cells >97%

Neutrophils were suspended in IMDM with 20% FBS and antibiotics at 1.11 x 10^6^ cells/ml and 225 μl was plated into each well of a 96 well plate. Monocytes were suspended in IMDM with 10% FBS and antibiotics at 5 x 10^5^ cells/ml and 1 ml was plated into each well of a 12-well plate. Cells were treated with *C. albicans* derived glucan (4 μg/ml), Pam3CSK4 (10 μg/ml, Invivogen), ultra-pure LPS (0.1 μg/ml, Invivogen) or a combination of Pam + LPS in duplicate for 18 h at 37C. PBS was used as the diluent control. A subset of cells was harvested to determine viability (>80%). The remainder were stained for SR-A expression. Each treatment was performed in duplicate, and each blood sample was run as an independent experiment.

### Flow Cytometry for Murine Cells

Flow cytometry for SR-A was performed on blood, bone marrow, splenic, and peritoneal leukocytes at 3, 16, or 24 as well as 2 or 7 d post-operatively. Whole blood was collected into sodium citrate. Splenocytes were isolated by teasing the tissue apart to liberate the cells and the stroma was discarded. Bone marrow as obtained by lavage of the femoral medullary cavity. Erythrocytes were lysed with PharmLyse buffer (BD Pharmingen). Cultured bone marrow cells or peritoneal cells were scraped into lidocaine. Leukocytes were blocked with 5% rabbit serum, 0.5% bovine serum albumin (BSA), and 5mM EDTA with anti-murine CD16/32 (BD Pharmingen) prior to staining. Leukocytes were stained with biotinylated rat anti-SR-A (clone 2F8, AbD Serotec, Oxford, UK), anti-Ly-6B.2 FITC (clone 7/4, AbD Serotec), and anti-F4/80 APC (AbD Serotec), or their isotype control antibodies. Staining was performed according to the protocol described by BD Pharmingen. Biotinylated antibodies were detected by streptavidin-phycoerythrin (BD Pharmingen). Cells were suspended in Pharmingen Stain Buffer and analyzed using a FACScalibur flow cytometer with CellQuest software (BD Biosciences, Mountain View, CA). LY6B.2 or F4/80 vs. side scatter plots were generated, and the neutrophil or monocyte/macrophage population was outlined, respectively (Figure S1). Histograms for SR-A were generated using the neutrophil or monocyte/ macrophage population as a gate and mean SR-A fluorescence was determined (Figure S1).

### Flow Cytometry for Human Cells

Whole blood RBCs were lysed with Pharmlyse (BD Biosciences, Mountain View, CA) according to package directions. Cultured neutrophils were lifted by gentle pipetting, and cultured monocytes were scraped into EDTA. Leukocytes were blocked with 5% rabbit serum, 0.5% bovine serum albumin, and 5 mM EDTA prior to staining. Leukocytes were stained with anti-SR-A APC (clone 351615, R&D Systems), anti-CD16 Pacific Blue (clone 3G8, BD Pharmingen), anti-CD14 FITC (clone TUK4, Bio-Rad Antibodies), or isotype control antibodies. Staining was performed according to the protocol described by BD Biosciences. Cells were fixed in 2% paraformaldehyde and stored in Pharmingen Stain buffer until analyzed. Cells were analyzed using a BD LSR Fortessa with FACSDiva software. Neutrophil and monocyte receptor mean fluorescences were determined by gating on the neutrophil population in a CD16 vs. side scatter plot (Figure S2) or the monocyte population in a CD14 vs. side scatter plot (Figure S3)

### Immunohistochemistry

Immunohistochemistry (ihc) for SR-A was performed on the lungs and spleens of CLP, sham, and control animals sacrificed at two days (n=4). Tissues were fixed in formalin and processed through paraffin. Lungs were not inflated. After treatment with a solution of blocking proteins followed by proteinase K treatment and then treatment with avidin and biotin, 6 micron paraffin embedded sections were incubated with biotinylated rat anti-mouse SR-A antibody (2F8, Serotec) diluted 1:200 for 3 h at 37 degrees. Biotinylated IgG of the same isotype as the primary was used as a control at 1:50. After washing, the sections were incubated in alkaline phosphatase conjugated streptavidin (Biogenex) for 15 minutes at room temperature. Warp Red was used as the chromogen according to the manufacturer’s instructions. Hematoxylin was used to counterstain and the sections were dehydrated and cover slipped. Images were analyzed by a pathologist blinded to conditions.

### Statistics

Graphing and statistical analyses were performed using GraphPad Prism 10. For the murine *in vivo* studies each time interval the flow data were normalized to the untreated control group which was assigned a mean fluorescence value of 100 (Figure 1, dashed line). Data is plotted as mean and standard error of the mean. Mean values at each time point were analyzed by analysis of variance, and experimental group means were compared with Tukey’s procedure. Sepsis clinical data were summarized by median and percentiles. For comparison of the two groups, the median mean fluorescence for each group was compared by Mann-Whitney Rank Sum. Correlations were determined by Simple Linear Regression. The SR-A mean fluorescence for the replicates (3 for murine, 2 for human) of the experimental groups in each experiment (4 for murine and 9 for human) was averaged and presented as the fold change from control (Figure 4, dashed line). Means (murine) or medians (human) were analyzed by analysis of variance, and experimental group means were compared with Tukey’s or Dunn’s procedure. Probability levels of 0.05 or smaller were used to indicate statistical significance.

**Figure 1.**
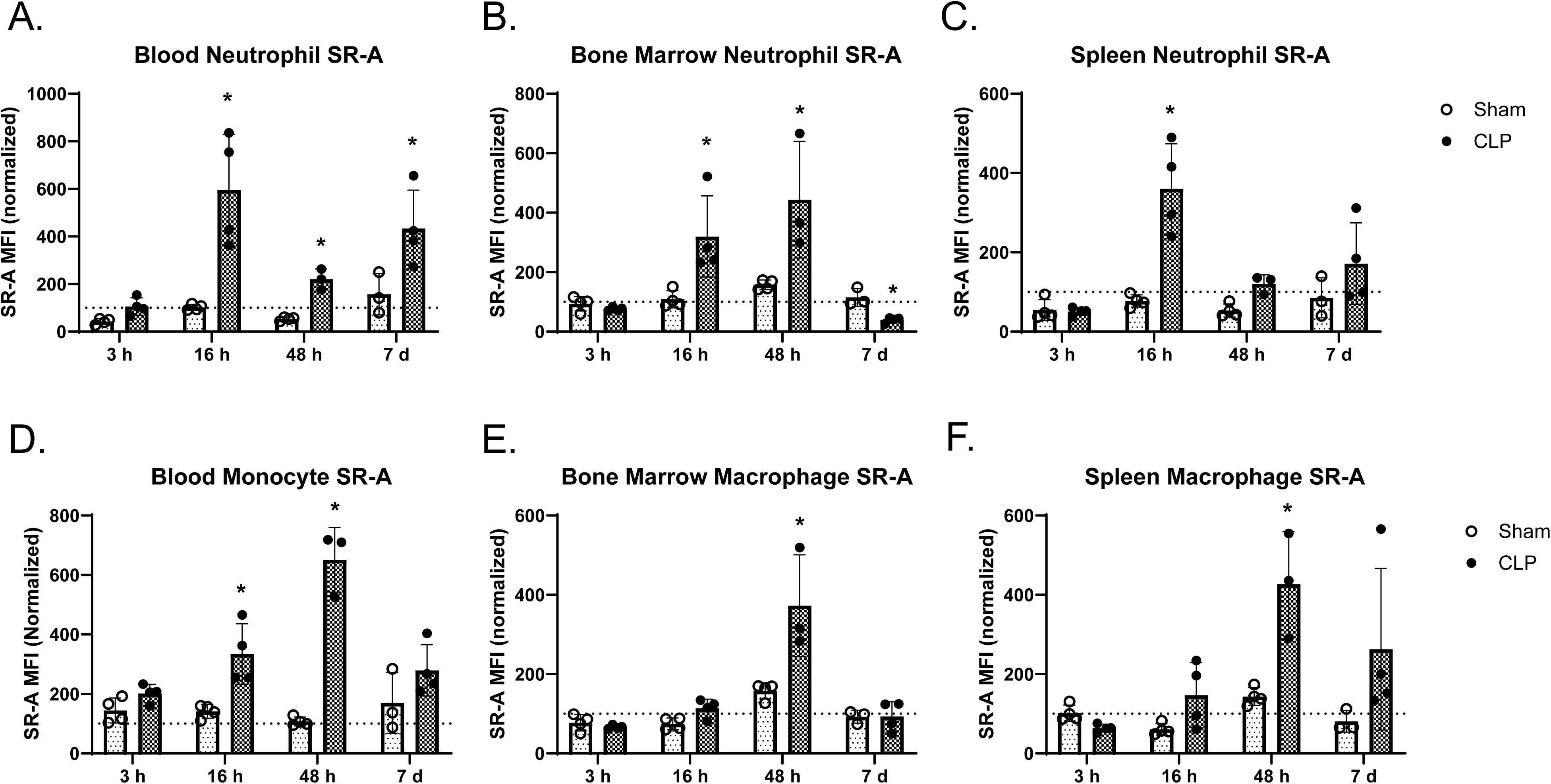
Polymicrobial sepsis increases SR-A expression on murine leukocytes. CLP or sham surgery mice were sacrificed at the indicated times. Blood (**A** and **D**), bone marrow (**B** and **E**), and spleens (**C** and **F**) were harvested, reduced to single cell suspensions, stained for cell type and SR-A, and analyzed by flow cytometry. SR-A mean fluorescent intensity was determined on neutrophils (**A-C**) by gating the population in an Ly6B2 vs. SSC plot an on monocytes/macrophages (**D-E**) gated in an F4/80 vs. SSC plot. Data presented have been normalized to control (dashed line). * Indicates p<0.05 compared to control and sham. n=3-4.

## Results

### Leukocyte SR-A is increased in a murine model of polymicrobial sepsis

Our previous study demonstrated that SR-A depletion improved survival in CLP sepsis (31). The improved survival was associated with decreased inflammation and decreased bacterial burdens. The goal of this experiment was to determine if sepsis alters myeloid cell SR-A expression. We found that blood neutrophils showed a significant increase in surface SR-A beginning at 16 h after CLP, and this increase continued throughout the course of acute sepsis and was maintained in those mice that survived at 7 d. Mean SR-A fluorescence was increased by 495% at 16 hr when compared to control and 482% compared to the sham surgery group (Figure 1A, p<0.05). Two days post-CLP SR-A expression was increased by 107% and 213% compared to control and sham, respectively (Figures 1A, p<0.05). In mice surviving to 7 d, SR-A fluorescence continued to be increased by 333% compared to control and 176% compared to sham operated mice (Figure 1A, p<0.05). Similar changes in cell surface SR-A expression were found in bone marrow neutrophils, though the kinetics of the change were somewhat different. SR-A mean florescence was increased in sepsis at 16 h (167% and 172% compared to control and sham, respectively, p<0.05) (Figure 1B). The increased SR-A expression peaked at 2 d (304% and 184% compared to control and sham, respectively, p<0.05) (Figure 1B). Interestingly, 7 d post-operatively, in the four surviving mice remaining from an initial ten surgeries, the mean fluorescence of SR-A on bone marrow neutrophils was decreased. When septic neutrophils were compared to control, SR-A was decreased by 60.6%, and when compared to sham surgery, SR-A was decreased by 65.7% (Figure 1B, p<0.05). In the spleen neutrophils showed increased SR-A expression at 16 h post-operatively (294% and 328% compared to control and sham, respectively, p<0.05). There was an increased SR-A expression at 2 d, this increase was only significant when compared to sham operated animals (126%, p<0.05) (Figure 1C). There was no significant difference in SR-A fluorescence between septic and non-septic splenic neutrophils 7 d after CLP (Figure 1C).

SR-A is classically considered a macrophage receptor (13–16). We sought to determine if sepsis alters SR-A expression on monocyte/macrophages. We found that sepsis increased SR-A expression on blood monocytes at 16 hours and 2 days post-operatively (Figure 1D). AT 16 h SR-A mean fluorescence on CLP monocytes was 381% increased over control and 460% increased over sham operated animals (Figure 1D, p<0.05). Circulating monocyte SR-A expression returned to control levels by 7 d post-CLP (Figure 1D). Bone marrow monocyte and macrophage SR-A was also increased in sepsis, but only at 2 d post-operatively (Figure 1E). SR-A mean fluorescence was increased on septic cells by 273% compared to control and 136% compared to sham at 2 d post-CLP (Figure 1E). Bone marrow monocyte/macrophage SR-A expression was not significantly different when comparing septic, sham, or control cells 7 d post-operatively (Figure 1E). Splenic macrophages from septic animals also had an increased mean SR-A fluorescence 2 d post-operatively (Figure 1F). SR-A expression on septic splenic macrophages was increased by 327% when compared to control and 197% when compared to sham macrophages (Figure 1F, p<0.05).

### Sepsis increases SR-A positivity in mouse lung and spleen

Sepsis is known to result in an infiltration of inflammatory cells, such as neutrophils, into the lung (7). These cells are thought to contribute to acute lung injury, and since our previous studies have shown that SR-A is detrimental in sepsis (31), we sought to determine if the pulmonary infiltrating cells express SR-A. Lung tissues were harvested at 2 d post-operatively and stained for SR-A expression by immunohistochemistry. After CLP, lungs showed an infiltration of SR-A positive cells (Figure 2A). Higher magnification revealed that many of these cells were consistent with macrophages, but smaller cells with less evident cytoplasm and segmented (neutrophil) or “u” shaped nuclei (“band” neutrophils), respectively, became quite numerous and evident at this time point (Figure S4A and B). The staining was cytoplasmic. Not all neutrophils or bands stained positively for SR-A (Figure S4A and S4B). We found that lungs from control mice had few SR-A positive cells morphologically consistent with alveolar macrophages (Figure 2A). The lungs of sham operated animals appeared similar (data not shown). These data support the conclusion that SR-A positive neutrophils, band cells and macrophages infiltrate the lung during sepsis.

**Figure 2.**
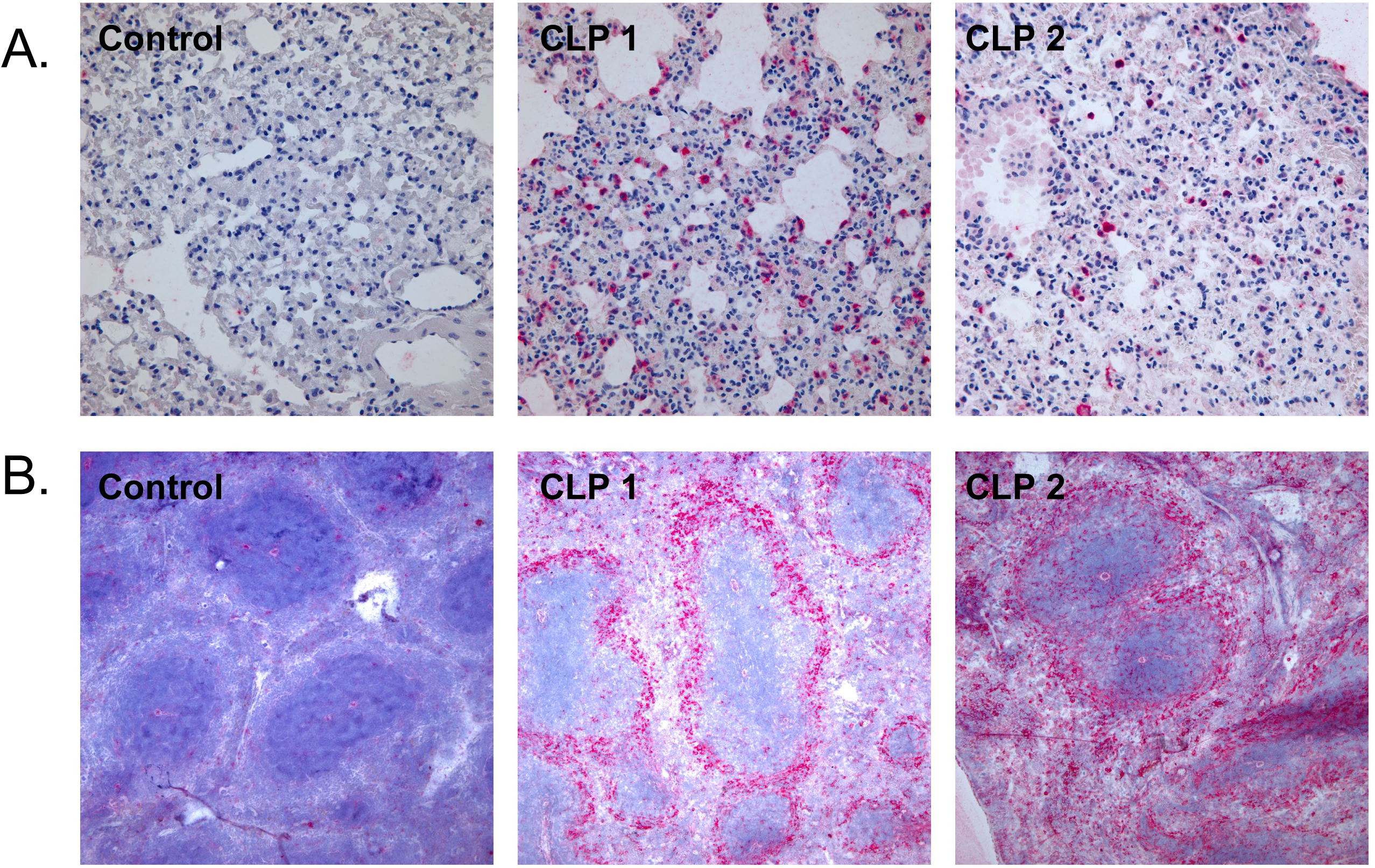
Polymicrobial sepsis increases SR-A expression in murine tissues. SR-A immunohistochemistry was evaluated in FFPE sections from the lungs (**A**) and spleens (**B**) of control and septic mice 2 d post-operatively. SR-A+ cells are stained red. **A.** While a few SR-A positive cells were observed in the lungs of control animals, sepsis induced a dramatic increase in SR-A positive pulmonary myeloid cells (red). **B**. There was a marked upregulation of SR-A on macrophages in the marginal zone surrounding the splenic nodules (red). Magnification 400x. Representative images from one control animal and two CLP animals.

Spleens were harvested from controls and at 2 d post-CLP and stained for SR-A expression by immunohistochemistry. Spleens from control animals show limited immunoreactivity in scattered cells appreciable only at higher power (Figure 2B). Spleens from CLP animals show pronounced immunoreactivity in the marginal area around splenic nodules (Figure 2B). The marginal area is populated predominantly by macrophages, and their immunoreactivity for SR-A has been markedly upregulated (red, Figure 2B and S4C). These data confirm those obtained by flow cytometry.

Taken together, these data demonstrate that in murine sepsis both neutrophil and monocyte/macrophage SR-A expression is increased in multiple tissues.

### Peripheral blood myeloid cell SR-A is not altered in sepsis patients

It is now well accepted that murine models of sepsis do not fully mimic the pathophysiology of sepsis (ref). Thus, we sought to determine if SR-A expression was also increased in sepsis patients. Patients were identified, consented, and enrolled by the ETSU Health Infectious Disease service at Johnson City Medical Center. Healthy control subject blood was purchased from a local vendor (BioIVT). The demographics of the subjects enrolled are described in **Table 1**. Fresh blood samples were stained for cellular phenotype and SR-A expression, and the mean fluorescence of SR-A on neutrophils and monocytes was determined by flow cytometry. We found that SR-A expression on circulating neutrophils and monocytes varied widely not only in sepsis patients but also in the baseline expression of healthy control cells (Figure 3A). When comparing the median surface expression, sepsis had no significant effect on SR-A expression in either neutrophils (p=0.585) or monocytes (p=0.141) when compared to healthy control subjects (Figure 3A). We sought to determine if the expression of SR-A in sepsis patients correlated with severity of disease. SOFA II scores were used as a marker of disease severity (1). We found that neither circulating neutrophil nor monocyte SR-A expression in sepsis patients correlated with SOFA II scores (Figure 3B). For SOFA II vs. neutrophil SR-A expression the R^2^ = 0.027 and the p = 0.165 (Figure 3B). For SOFA II vs. monocyte SR-A expression the R^2^ = 0.025 and the p = 0.175 (Figure 3B). Thus, SR-A expression in septic leukocytes from patients does not correlate with SOFA II scores or disease severity.

**Figure 3.**
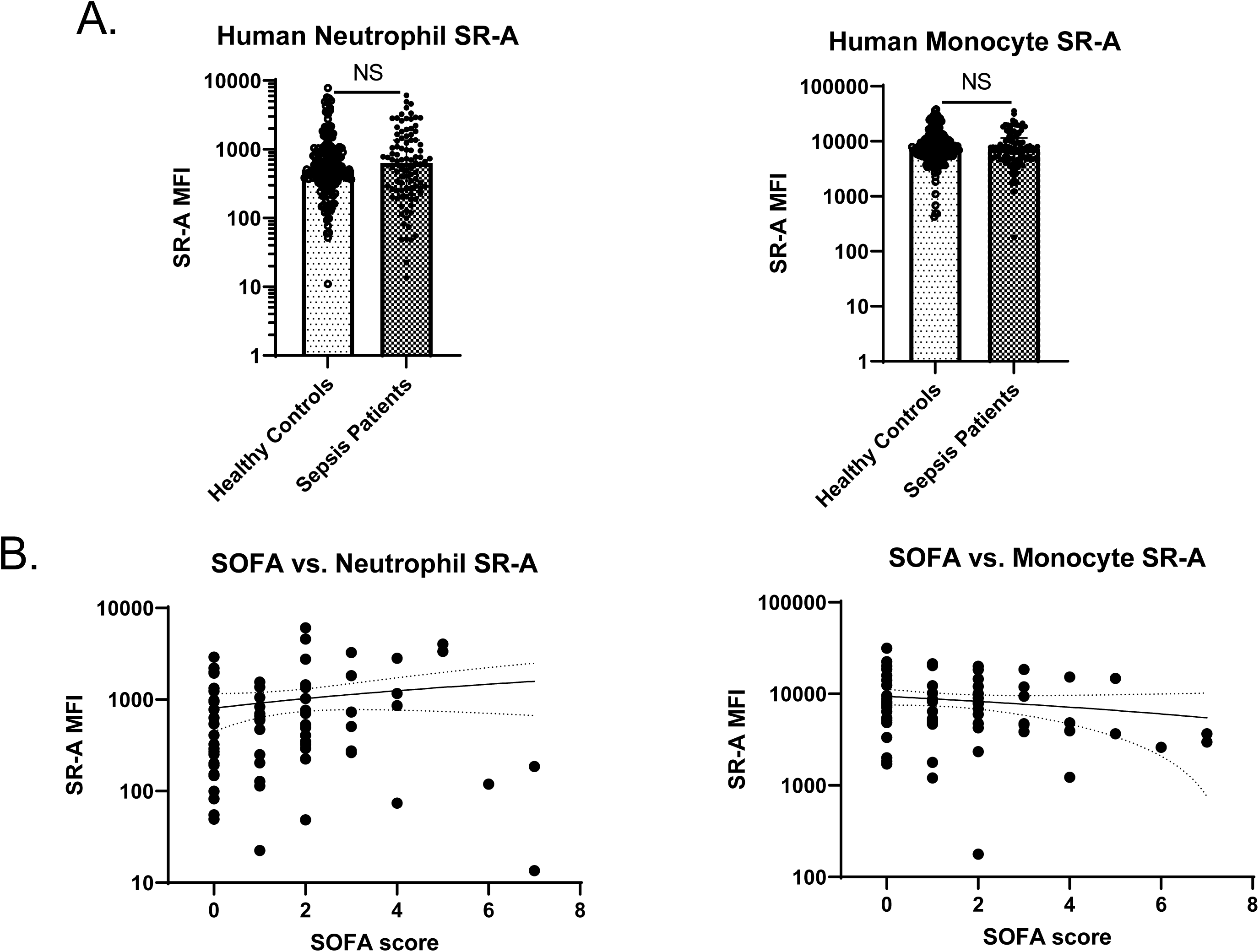
SR-A expression on blood leukocytes is not altered in sepsis patients, and SR-A expression does not correlate with disease severity. Leukocytes were harvested after informed consent and isolated by red cell lysis, followed bystaining for cell phenotype and SR-A expression. **A.** SR-A mean fluorescent intensity was determined for the neutrophil population after gating the population in a CD16 vs. SSC plot and for the monocyte population after gating the population in a CD14 vs. SSC plot. n = 105-181; NS indicates p>0.05. **B.** SOFA II scores were recorded and plotted against neutrophil SR-A MFI and monocyte MFI. Linear regression detected no correlation between SR-A expression and SOFA II scores. n = 78, p>0.05.

**Table 1.** Human subjects demographic data.

|  | **Healthy Controls<br>(N = 181)** | **Sepsis Patients<br>(N = 105)** |
|---|---|---|
| Mean Age yrs (range) | 53.6 (30-73) | 60.6 (24-95) |
| % Female | 52% | 47% |
| Race (%): White | 91.7% | 100% |
| African American/Black | 5.5% | 0% |
| Latino/a/x | 0.5% | 0% |
| Asian | 2.2% | 0% |

### PAMP stimuli increase murine leukocyte SR-A, but not human leukocyte SR-A

Our data demonstrate that murine and human leukocytes show differential SR-A responses in sepsis. We sought to determine if the leukocyte SR-A responses were specific to sepsis. To this end, we treated primary murine bone marrow neutrophils or elicited peritoneal macrophages with ultrapure LPS, a TLR4 ligand, Pam3CSK4, a TLR2 ligand, *C. albicans* β-glucan, a Dectin-1 ligand, a combination of LPS and Pam3CSK4 (TLR2 and TLR4) or media alone for 24 hours. Cell type and SR-A mean fluorescence were determined by flow cytometry. In the murine cells we found that only the TLR2 ligand Pam3CSK4 significantly altered SR-A expression in bone marrow neutrophils (Figure 4A). In bone marrow neutrophils Pam3CSK4 increased SR-A expression by 569% (p<0.05; Figure 4A). LPS alone did not significantly increase SR-A expression, but Pam3CSK4 in combination with LPS increased SR-A expression on bone marrow neutrophils by 573% (p<0.05; Figure 4A). β-glucan also had no significant effect on SR-A expression on bone marrow neutrophils (Figure 4A). Thus, only TLR2 stimulation mediates increased murine neutrophil SR-A expression, while neither TLR4 nor Dectin-1 stimulation have an effect on SR-A expression (Figure 4A). In contrast, mouse peritoneal macrophage SR-A expression was significantly increased not only by TLR2 stimulation, but also by TLR4 stimulation, though the degree of increase was not as high as that seen in TLR2 stimulated mouse neutrophils (Figure 4B). Specifically, LPS alone increased SR-A expression on macrophages by 68.2% while Pam3CSK4 alone increased SR-A expression by 46.1% when compared to control cells (p<0.05; Figure 4B). Interestingly, the combination of LPS with Pam3CSK4 did not significantly increase SR-A expression above that with either mediator alone, though it was still significantly above that of the controls (p<0.05; Figure 4B). Much like with neutrophils, glucan treatment did not significantly alter mouse macrophage SR-A expression (Figure 4B).

**Figure 4.**
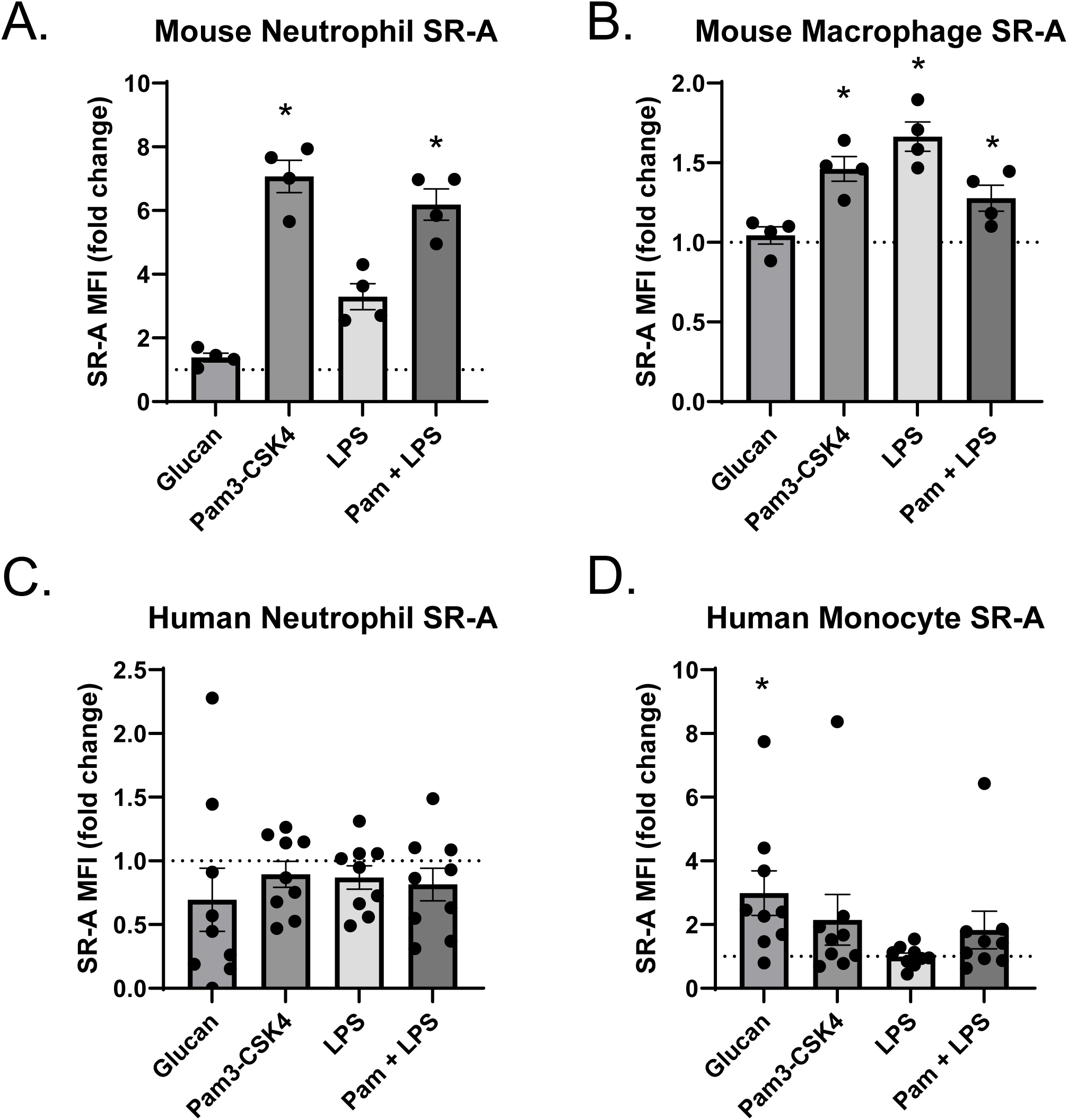
PAMP stimulation increases SR-A expression on murine leukocytes, but does not change SR-A expression on human leukocytes. **A**. Murine bone marrow neutrophils were incubated for 24 h with mediators followed by staining and flow analysis for SR-A MFI. Data points are four independent experiments run in triplicate and presented as fold change from control (dashed line). * indicates p < 0.05 compared to untreated control. **B**. Murine thioglycolate elicited macrophages were incubated with mediators for 24 h followed by staining and flow analysis for SR-A MFI. Data points are four independent experiments run in triplicate and presented as fold change from control (dashed line). * indicates p < 0.05 compared to untreated control. **C**. Human neutrophils were incubated with mediators for 18 h followed by staining and flow analysis for SR-A MFI. Data points are from 9 independent experiments run in duplicate and presented as fold change from control (dashed line). P>0.05. **D**. Human monocytes were incubated with mediators for 18 h followed by staining and flow analysis for SR-A MFI. Data points are from 9 independent experiments run in duplicate and presented as fold change from control (dashed line). * Indicates p < 0.05 compared to control.

In stark contrast to murine neutrophils, treatment of human blood neutrophils with the same mediators for 18 h induced no significant change in SR-A expression compared to untreated control (Figure 4C). When blood monocytes were treated with the stimuli, we found that only glucan treatment resulted in a significantly increased SR-A expression compared to untreated controls (Figure 4D). There was a 116% increase in SR-A mean fluorescence with glucan treatment than PBS alone (p<0.05). This is also in contrast to the response of murine macrophages when treated with these stimuli in which all PAMPs except glucan had an effect.

## Discussion

We have reported that SR-A mediates, in part, the pathophysiology of sepsis in a murine model of polymicrobial sepsis (31). The current study extends our findings on SR-A in CLP sepsis by demonstrating that septic mice show increased expression of SR-A on myeloid cells both in circulation and in the tissues compared to sham operated and control animals, thus supporting the role of SR-A in mediating the detrimental effects of CLP sepsis. In striking contrast, leukocytes from sepsis patients did not show SR-A up regulation. Not only was myeloid cell SR-A unchanged in sepsis, but SR-A expression did not correlate with disease severity. Additionally, SR-A expression was not affected by PAMP treatment of human neutrophils or monocytes. In contrast, murine cells showed increased SR-A expression in a TLR2 (neutrophil) or TLR2/TLR4 (macrophage) dependent manner. These data indicate that with respect to leukocyte SR-A there is a differential response between humans and mice in sepsis and with respect to pathogen associated molecular patterns.

Though we believe that the discrepancy between the SR-A response in sepsis patients and septic mice is due to differences in the regulation of SR-A expression, it is possible that the differences could be experimental. The CLP model used here is more severe compared to that of the sepsis patients. The majority of sepsis patients had mild to moderate disease, with only a few considered severe sepsis and no mortalities, whereas this model of CLP generally has a mortality rate of 60-80% (31 and data not shown). That being said, the patients on the higher end of the severity scale tended to have similar or lower neutrophil and monocyte SR-A expression compared to mild or moderate cases. Sepsis is a heterogenous disease with multiple inciting organisms, inciting sites of infection and clinical presentations (1). Our mouse model is a polymicrobial infection of the peritoneal cavity that advances to sepsis, while the majority of enrolled sepsis patients had different single organism infections with varying sites of infection including pneumonia, urinary tract infection, and cellulitis. Variability in infectious agents could certainly account for the differences in the SR-A response.

However, the *in vitro* studies suggest that the difference is due to differences in SR-A transcriptional and/or translational regulation of SR-A expression between the two species in response to infectious stimuli. Though human myeloid cells express PRRs and respond to these PAMPs in other ways, they do not respond by upregulating SR-A, unlike murine myeloid cells. This suggests a differential regulation of SR-A in response to PAMP stimulation in humans versus mice. The mechanisms responsible for changes in SR-A expression under different conditions have been little studied (25). Further, there seems to be little delineation between studies performed in human cells and mouse cells (25). Additional studies are needed to compare regulation of SR-A expression in human cells in response to different condition to that in mouse cells.

Though the alterations in SR-A with CLP sepsis in mice were not recapitulated in sepsis patients, it is possible that the overall effect that SR-A has on cellular function could relate to pathophysiology of sepsis on a broader scale. For instance, SR-A stimulation in neutrophils has been associated with increased NET production (18,19). NETs are integral to the neutrophil response to infection, but they contain multiple inflammatory damage associated molecular patterns (DAMPs) including histones and extracellular DNA. NETs also tend to be pro-coagulation (8,35,36). In sepsis, neutrophils have dysfunctional migration, and NETosis will occur in the microvasculature (5,6,37). The increased inflammation and clotting associated with NETs in the microvasculature has been associated with organ damage in sepsis (8,35,36), therefore an increase in NETosis due to increased neutrophil SR-A expression could lead to poorer survival in septic wild type animals compared to SR-A knockouts. Alternatively, increased SR-A expression has been found on immunosuppressive tumor associated macrophages, also known as M2 macrophages (38,39). An increase in SR-A expression on macrophages in septic mice would suggest an increase in immunosuppressive macrophages which would impair immune response to ongoing infection. Additionally, in dendritic cells it has been shown that SR-A expression suppresses T cell activation (40). We did not specifically examine dendritic cell SR-A, but it is possible that SR-A expression has a similar effect on macrophage/T cell interaction leading to suppression of both innate and adaptive immune responses to ongoing infection. Many researchers believe that immunosuppression is the primary cause of morbidity and mortality in sepsis patients (9). Thus, an increase in immunosuppressive macrophages could be responsible for the increased mortality and bacterial burdens in wild type septic mice compared to SR-A deficient. Though the mouse CLP model of sepsis does not reflect the role of SR-A in septic patients, the overreaching changes in cellular mechanisms due to the increased SR-A in mice could potentially translate.

The data presented show that myeloid SR-A expression is increased in murine sepsis, and based on previous data increased SR-A plays a detrimental role in the response to sepsis (31). In contrast, SR-A expression is not changed in leukocytes derived from septic patients and does not correlate with SOFA II scores. Thus, myeloid SR-A expression does not appear to play significant a role in the pathophysiology of human sepsis.

## Authorship contribution

TRO: experimental design, data acquisition (CLP surgeries, human and murine cell isolation, culture, and treatment, flow staining and acquisition), data analysis, original draft and revision;

MDM: data acquisition (human cell isolation and culture, flow acquisition), data analysis, review, editing;

JA: clinical coordinator;

NH: data acquisition (murine cell isolation and culture, flow acquisition);

JBS: data acquisition (pathology); and DLW: project oversight, source of reagents, review, editing and revision

---

**Figure S1.**
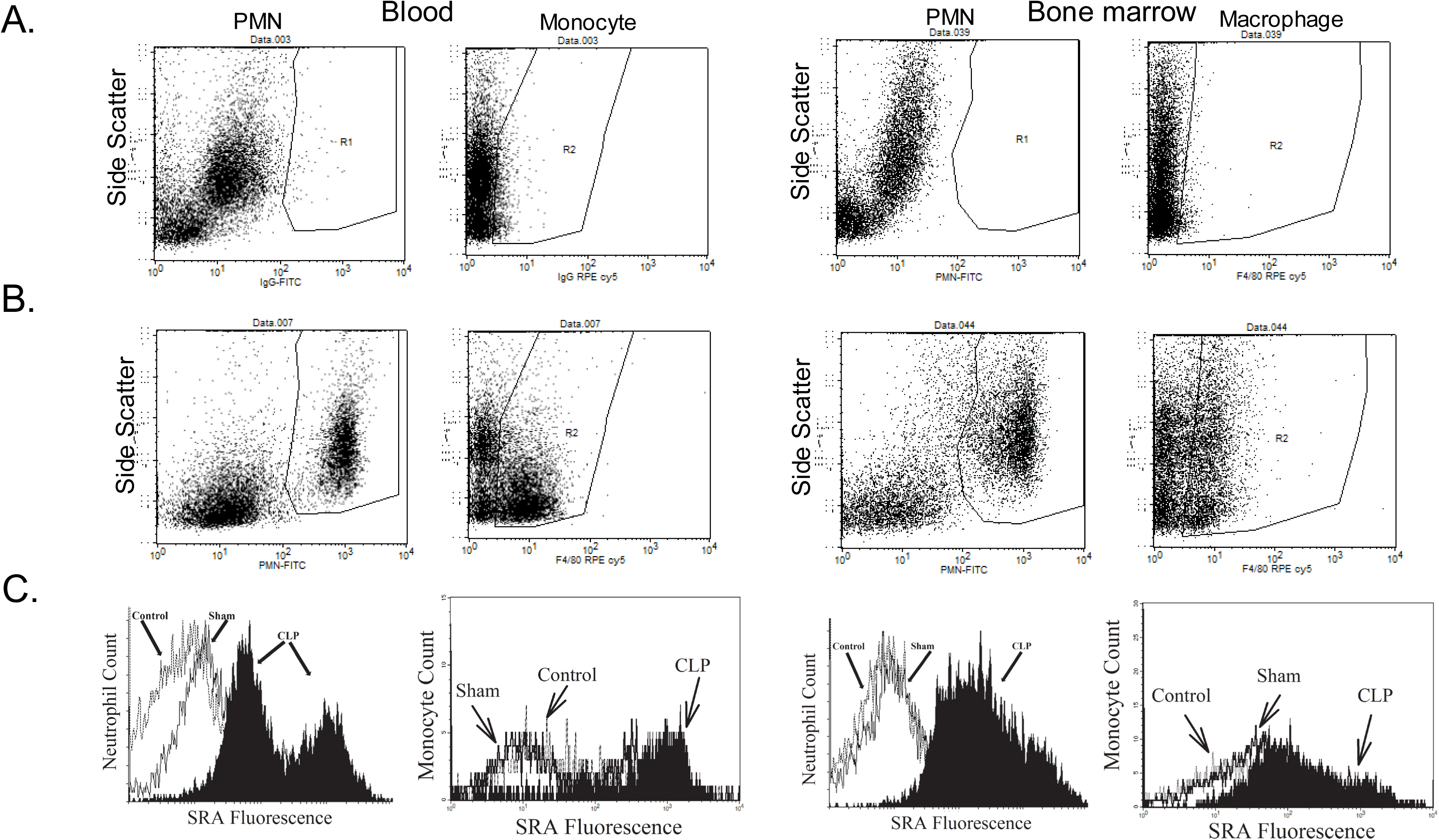
Gating strategy and raw data for murine flow cytometry. Blood or bone marrow leukocytes were stained with either isotype control antibodies or Ly6B2-FITC, F4/80-PE, and SR-A-APC. **A**. Gates for the cell type were established using the isotype stained cells in FITC vs. SSC (neutrophils) or PE vs. SSC (monocyte/macrophage) plots. Cells from the neutrophil or monocyte/ macrophage gate (**B)**. were used to generate histograms to determine SR-A mean fluorescent intensity (**C)**.

**Figure S2.**
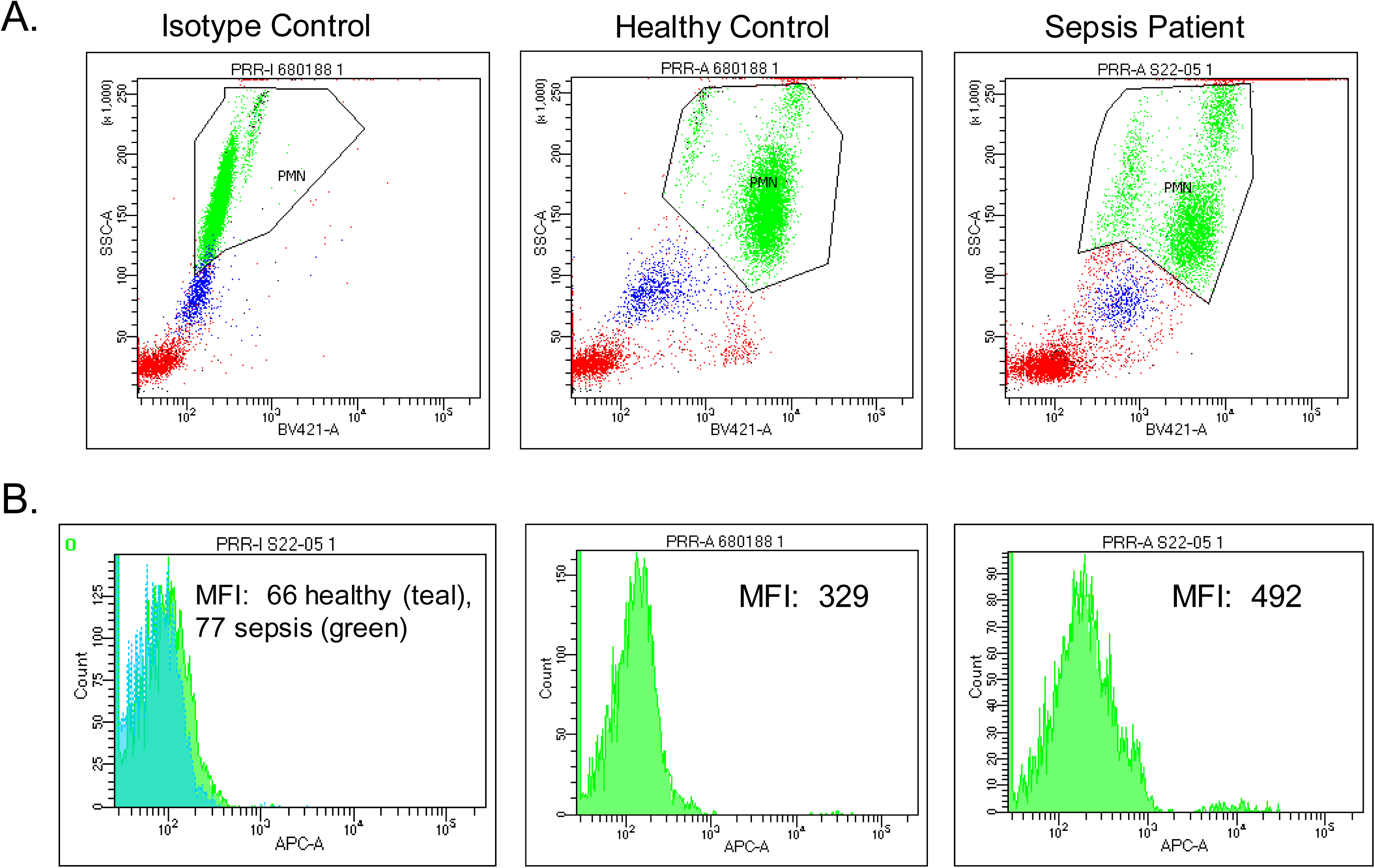
Gating strategy for human neutrophil flow cytometry. Blood leukocytes were stained with either isotype control antibodies or CD16 Pacific Blue and SR-A APC. **A**. Neutrophils were gated in a CD16 vs. SSC plot. **B**. APC mean fluorescence was determined on the cells from the neutrophil gate. Representative histograms of isotype control (healthy in teal, sepsis in green), healthy control, and sepsis patient.

**Figure S3.**
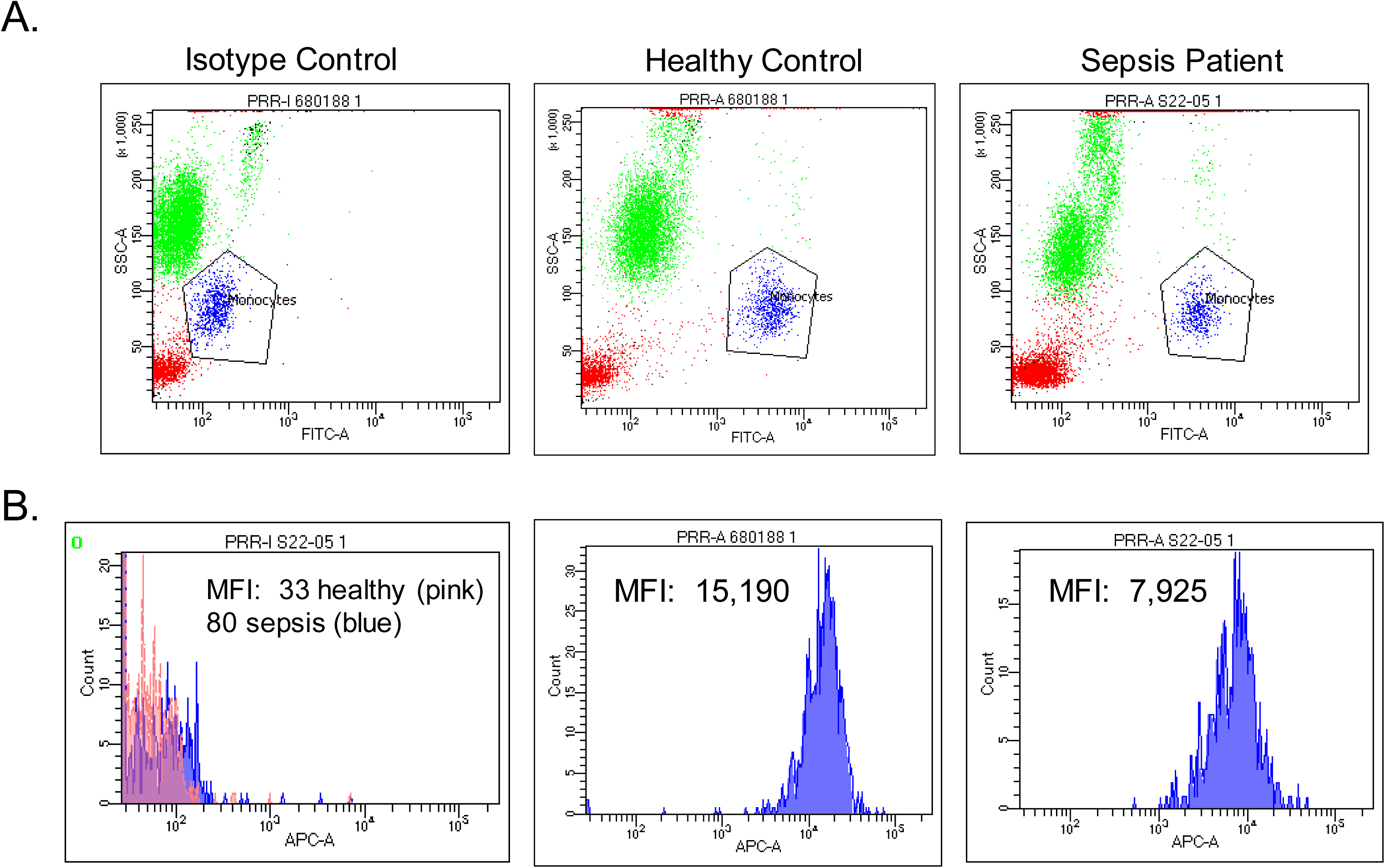
Gating strategy for human monocyte flow cytometry. Blood leukocytes were stained with either isotype control antibodies or CD14 FITC and SR-A APC. **A**. Monocytes were gated in a CD14 vs. SSC plot. **B**. APC mean fluorescence was determined on the cells from the monocyte gate. Representative histograms of isotype control (healthy in pink, sepsis in blue), healthy control, and sepsis patient.

**Figure S4.**
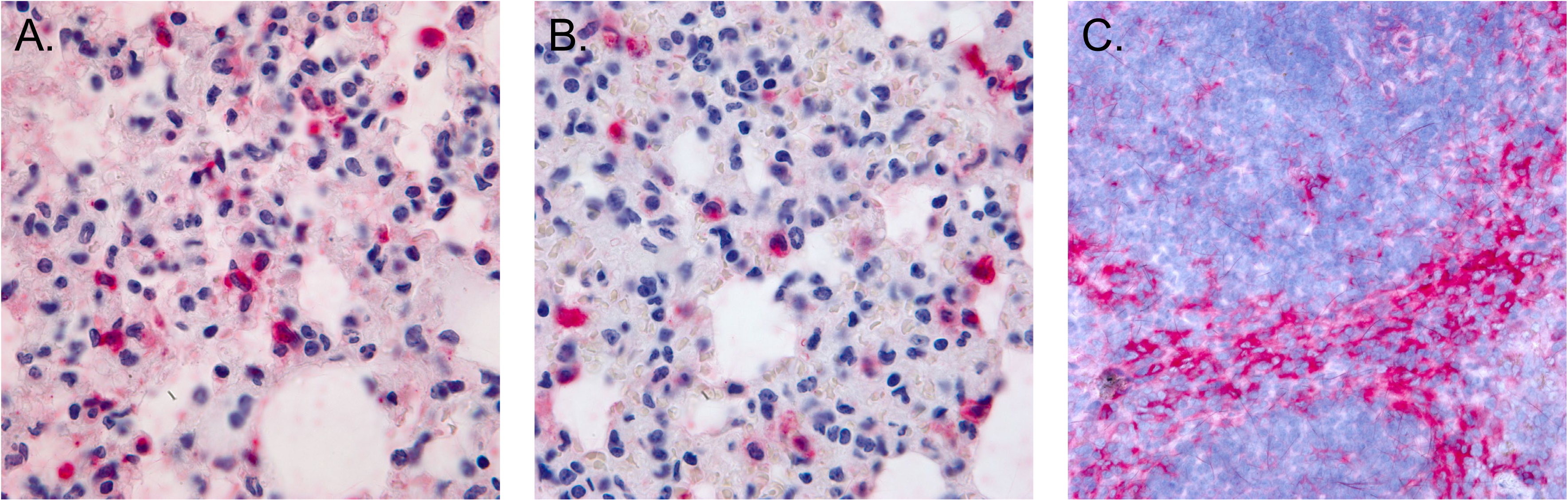
Higher power images of septic lung (**A** and **B**) and spleen (**C**) to show cellular details. SR-A positive cells (red) include cells with monocyte, immature band neutrophil, and mature neutrophil morphology in the lung, though there are cells of these types that are negative for SR-A (**A** and **B**). SR-A positive cells in the spleen are primarily macrophages (**C**). Magnification 1000X.

